# Tracking niche change through time: simultaneous inference of ecological niche evolution and estimation of contemporary niches

**DOI:** 10.1101/2019.12.29.890608

**Authors:** Xia Hua, Marcel Cardillo, Lindell Bromham

## Abstract

1. The ecological niche of species is often studied from two different perspectives. Environmental niche modelling (ENM) infers a species’ current niche from contemporary occurrence records, while phylogenetic comparative methods (PCM) infer the history of niche evolution. Although these two areas of research are conceptually linked, they are analysed independently, within separate analytical frameworks.
2. Here we provide a new method, NEMo (Niche Evolution Model), for simultaneous inference of niche evolution and estimation of contemporary niches of species. NEMo explicitly models three fundamental processes of niche evolution – adaptation, speciation, and dispersal – applying a reversible jump algorithm to infer occurrences of these processes on a phylogeny. The model permits ENMs to account for the role of history in shaping current species distributions, and offers more realistic models of evolution in PCMs.
3. Simulations show that NEMo has high accuracy for estimating the ecological niche of species, and reasonable power to identify the occurrences of the three processes on phylogeny. When applied to a real case study (the Australian plant genus *Acacia*), the method is more effective at predicting a key environmental niche axis (salt tolerance) than using ENM alone, and it infers temporal patterns in the evolution of drought tolerance in response to aridification that are consistent with prior expectations
4. NEMo makes it possible to combine many types of data to study niche evolution and estimate species niches, not only species distributions and phylogeny, but also paleoclimate, species tolerance range, and fossil records.

## Introduction

There is a plethora of different ways of defining and describing ecological niches. Here we focus on the Hutchinsonian niche (Hutchinson, 1957), the range of environmental conditions in which a species can establish or persist (Holt, 2009). This can be described as a mathematical distribution along one or more salient environmental axes, which can be estimated from species distribution and environmental data using ecological niche models (ENM) (Peterson et al., 2011). Ecologists often have strong motivations for estimating a species’ ecological niche. One is to define the range of conditions under which the species can persist: this information is useful for current conservation planning (e.g., Weber et al., 2017) or for predicting future distributions given a change in environmental conditions (e.g., González-Orozco et al., 2016). In addition, we may be interested in reconstructing the history of change in a species niche over time, to describe the way lineages respond to environmental change (e.g., Cang et al., 2016) or diversify to occupy a range of conditions (e.g., Castro-Insua et al., 2018). Typically, these two areas of inference – modelling species present-day niches and reconstructing niche evolution over time – have been conducted separately. But these two areas are conceptually linked, because the current niches of species are the product of evolutionary history, and reconstructions of niche evolution usually work backwards from data on current species’ niches. Estimating contemporary niches and inferring niche evolution separately has a number of shortcomings, explained in detail below.

ENM estimates a species ecological niche as a mathematical representation of the environmental conditions in the places where the species is known to occur or not occur (Peterson, 2011). Correlative ENM makes the strong assumption that a species current distribution is determined by its current ecological niche (Dormann et al., 2012), but this assumption is not always realistic. The current distribution reflects not only the environmental tolerances of the species, but other ecological constraints such as dispersal and interspecific competition (Saupe et al., 2018), as well as evolutionary history. Mechanistic ENM (Kearney & Porter, 2009), and more generally, process-based ENM (Dormann et al., 2012) can account for the ecological constraints on species current distribution, but not the historical role of speciation and adaptation in determining species’ present-day niches. While there have been several recent developments of ENMs that incorporate a phylogenetic dimension (Smith et al., 2018; Morales-Castilla et al., 2017; Maguire et al., 2018), these have mostly aimed to account for the statistical problem of non-independence due to relatedness among contemporary species, rather than to model the historical development of niches through evolutionary time.

The evolution of niches has previously been modelled in the framework of phylogenetic comparative methods (PCM), which model the evolution of biological traits in a group of species based on their current trait values and phylogenetic relationships (Felsenstein, 1985). PCMs have been applied to analyses of niche evolution by considering the ecological niche effectively as a species’ trait (Kozak & Wiens, 2010), which is often quantified as a point estimate (e.g., mean) of one or more environmental variables across species occurrence locations. The evolution of the niche can then be inferred along the phylogeny using PCMs. Modelling niches in this way presents three important problems. First, it has the same problem as ENMs that current species occurrence data alone may not give reliable estimates of a species’ ecological niche (Saupe et al., 2018). Second, a species’ niche is often described as a distribution along key environmental gradients, but PCMs were designed to work with point estimates rather than distributions (Hardy, 2006). Third, niche evolution involves complex, and still incompletely understood, biological processes, which may not be adequately characterized by the kinds of models (usually Brownian Motion or one of its variants) used to characterize trait evolution in PCM (Revell et al., 2008). Because of these limitations, models for trait evolution used in PCMs have been criticized for lacking biological realism (Münkemüller et al., 2015) and being hard to interpret.

We can overcome many of the limitations of these two alternative approaches – ENM and PCM - by combining the strengths of both in a common framework. To combine ENM and PCM, we take a Bayesian approach to simultaneously inferring niche evolution and estimating contemporary niches of species, with a novel process-based model (Niche Evolution Model: NEMo). NEMo is based on three biological processes expected to drive niche evolution and influence species distribution: adaptation, speciation, and dispersal. As for all modelling procedures, we need to consider a tradeoff between model complexity and model flexibility. NEMo cannot provide a comprehensive description of all aspects of the complex process of niche evolution, but by linking ENM and PCM, it increases model flexibility by incorporating biologically meaningful parameters that can be directly informed by various data types (see details in Discussion). In this paper, we present NEMo as a general framework to link ENM and PCM. It can be tailored to specific taxa, given its flexibility in incorporating our prior knowledge on the taxa and in linking different types of ENMs and PCMs.

## Methods

### Basic model structure

Figure 1 summarizes the basic structure of our approach. There are three major components: the phylogenetic comparative method (PCM), the model of niche evolution (NEMo), and the ecological niche model (ENM). We can express these three components in terms of conditional probability as follows. PCM estimates *p*(model|phylogeny), which is the probability of a certain history of niche evolution along a given time-scaled phylogeny. NEMo estimates *p*(niche|model, phylogeny), which is the probability of evolving the inferred contemporary niche of each tip species given the history of niche evolution and the phylogeny. ENM estimates *p*(data|niche), which is the probability of the distribution data of each tip species, given our inferred niche of the species. These probabilities provide the general framework of our method:

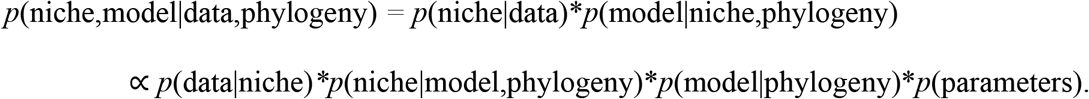

The inputs - (data, phylogeny) – will usually be distribution data with the environmental conditions of interest (such as the degree of aridity or salinity) at each presence location, and in some cases, reports of definite absence of the species from other locations, plus a time-scaled phylogeny of the taxa. The outputs or inferences - (niche, model) - are the posterior distribution of the ecological niche of each tip species and the posterior distribution of the history of niche evolution. Figure 1 summarizes the parameters to estimate in the Bayesian inference. We describe these parameters in each of the three components of our approach in the next three sections.

**Figure 1.**
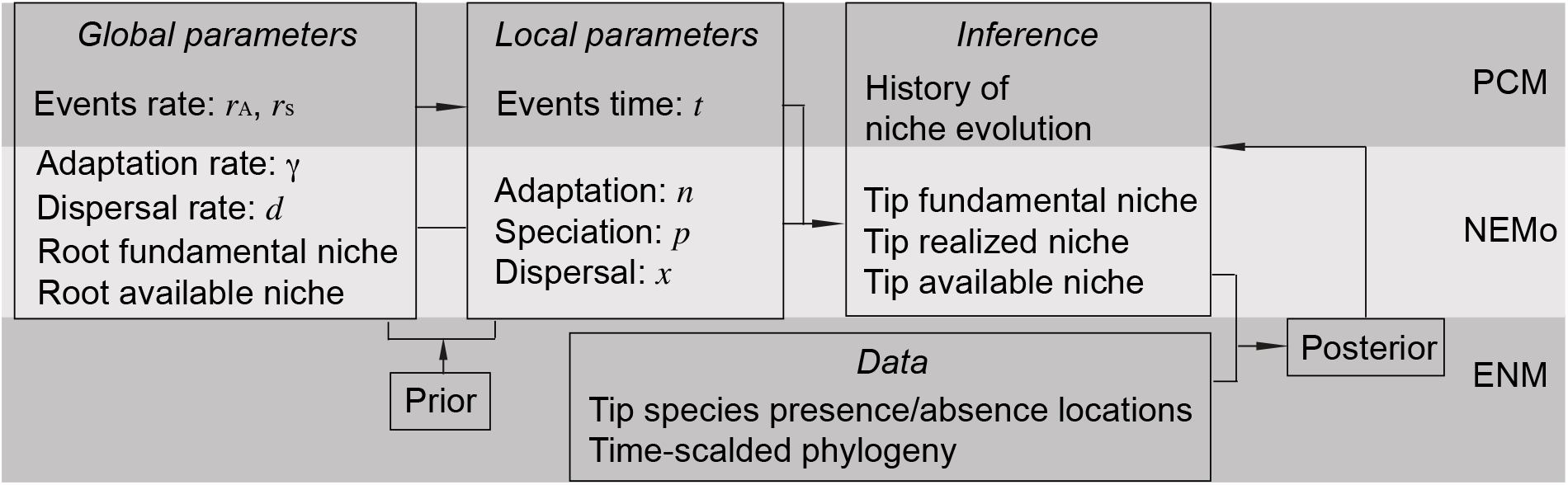
Illustration of the basic model structure. See text for details.

### NEMo: process-based niche evolution model

NEMo models three key biological processes that can shape niche evolution: adaptation, speciation, and dispersal. Adaptation and speciation are modelled as discrete events that occur at particular points on the phylogeny, and dispersal is modelled as a continuous process between two discrete events. Each process has distinct properties modelled by local and global parameters. Local parameters are those that are estimated for each event, such as the location of the event on the phylogeny (see the PCM section) and the way that a species’ niche changes during the event. Global parameters are those that are shared among events, such as adaptation rate and dispersal rate (Fig. 1).

The model starts from the ancestral species niche at the root of the phylogeny. At each internal node, two descendant species inherit niches from their most recent common ancestor, and their inherited niches may change at the node. Along each branch of the phylogeny, a species’ niche changes independently from other species, and the change depends on the order and the type of events occur along the branch. To make the model tractable, we only consider the expected change in species ecological niche under assumptions on the probability distribution of environmental conditions in an area and random species’ movement into areas with environmental conditions under which it can persist (see below), so a species’ ecological niche evolves in a deterministic way, i.e., *p*(niche|model, phylogeny)=1, and NEMo is not a spatially explicit model for niche evolution. In the Discussion, we discuss the idea of a joint stochastic process of both niche evolution and species movement over space. However, this is not computationally cost-effective and tractable when we don’t have detailed data to inform past species distributions, so we focus on the deterministic model here.

Specifically, the model describes expected changes in three aspects of a species niche (fundamental niche, available niche, realised niche) during adaptation, speciation, and dispersal processes (Fig. 1). Each of these aspects of the niche is described as a probability distribution along a particular environmental axis of interest for the system under study (e.g., drought or salinity), but can be extended to include multiple environmental factors. For the purposes of this model, we take the fundamental niche to be the probability that a species can persist under an environmental condition (a point along the environmental axis of interest).

We consider the available niche to be the relative frequency of different environmental conditions in the potential areas that the species can expand into and persist under the condition. The realised niche is taken to be the relative abundance of the species under each environmental condition. These niches follow the BAM conceptual framework of distributional ecology (Peterson et al., 2011), with the fundamental niche corresponding to the “A” space, the abiotically suitable area for a species; the available niche corresponding to the intersect of “M” and “B” space, the area accessible to the species via dispersal and not constrained by biotic interactions; the realized niche corresponding to the intersect of “A”, “M”, and “B” spaces, the area in which one can find the species.

To estimate the expected changes in the three aspects of a species niche, we assume that, for a key environmental axis in a particular study system, the spatial variation along this environmental axis in an area follows a truncated normal distribution, such that the distribution is truncated at dispersal barriers to the species (see details on dispersal). When this distribution shifts along the environmental axis, it induces selection on the species’ fundamental niche, and we assume the species’ fundamental niche evolves according to the Breeder’s equation, which results from differential survival and reproduction of individuals that can persist under different environmental conditions (see details on adaptation and speciation). We also assume that species randomly move over space with a dispersal rate *x*, which defines how rapidly the species will eventually occupy all environmental conditions under its available niche (see details on dispersal).

Our model starts from the three aspects of the ancestral species niche at the root of the phylogeny. Here, we focus on the evolution of tolerance to extreme conditions, so environmental conditions are more extreme towards the right of the axis (Fig. 2). The fundamental niche of the root species is assumed to follow *f*(*x*) ***=*** *e*^−*(x/α*)*β*^ on the environmental axis, so that under the least extreme condition, fitness *f*(0) =1. Parameters *α* and *β* control the scale and the shape of the niche: the larger the *α*, the more spread out the niche; the smaller the *β*, the slower the fitness declines towards extreme conditions (Fig. 2). Note that, by modifying the form of the environmental axis and the distribution of fitness under different conditions, our method can be adapted to questions other than the evolution of tolerance to extreme conditions. The available niche of the root species is assumed to follow a normal distribution on the environmental axis. The realized niche of the root species equals the product of its fundamental niche and its available niche: the distribution of the root species is at equilibrium, including all the areas of suitable habitat that the species is able to disperse to and occupy.

**Figure 2.**
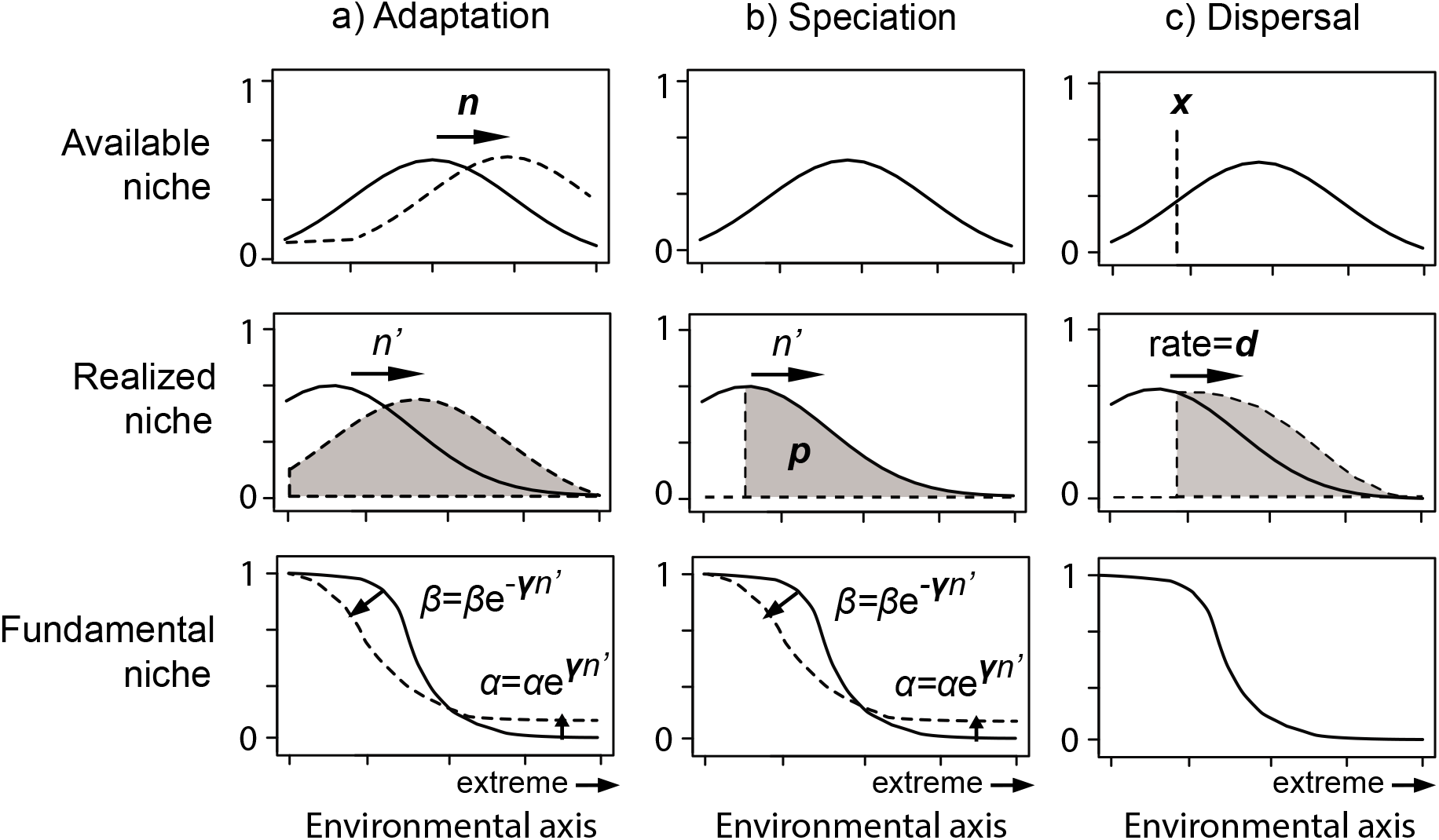
Illustration of the process-based model for niche evolution. There are three types of events: adaptation (a), speciation (b), and dispersal (c). Each type of event may change three aspects of niches along an environmental axis (with more extreme environmental conditions towards the right of the axis). In each plot, the solid curve is the niche before the event and the dashed curve is the niche after the event if any change happens (realized niche after the event is also shaded in grey). Bold letters are parameters to be estimated for the type of event. Each type of event is described by two parameters: dispersal rate (*d*) and the location of barrier on the environmental axis (*x*) for a dispersal event; adaptation rate (*γ*) and the amount of environmental change (*n*) for an adaptation event; adaptation rate (*γ*) and the location of species split on the environmental axis (*p*) for a speciation event. *n’* is the inferred difference in the mean of the realized niche before and after an event. Together with adaptation rate (*γ*), they determine change in fundamental niche through *α* and *β*, where increase in *α* leads to higher fitness under extreme condition and decrease in *β* leads to less abrupt decline in fitness towards extreme condition.

Now we can describe how these three aspects of a species niche change due to adaptation, speciation, or dispersal processes. An adaptation event (Fig. 2a) is triggered by a change in the environmental conditions in the species current distribution range. We model the environmental change as a shift by *n* towards extreme conditions in both the species’ realized niche and available niche (Fig. 2a). This is to capture an environmental change that not only affects the suitability of the species currently occupied area (realized niche) but also influences the suitability of other areas that it has the capacity to disperse to (available niche). As a result of differential survival of individuals under different environmental conditions (but before any dispersal occurs), the new realized niche is the product of the fundamental niche and the old realized niche shifted by *n*, which gives the relative abundance of individuals surviving in the area they used to occupy. As a result, the final amount of shift from the old realized niche to the new realized niche is not *n* and we denote it as *n*^*′*^ in Figure 2a. Treating each individual’s tolerance range as a trait, then *n*^*′*^ is the selection differential in the Breeder’s Equation, or the change in the mean tolerance range of the species over a generation if tolerance range has complete heritability.

Along with the change in the realized niche, the species fundamental niche can also evolve, resulting in a shift toward higher fitness in extreme conditions. According to our parameterization of the fundamental niche, higher *α* leads to higher fitness in extreme conditions and lower *β* leads to less abrupt decline in fitness towards extreme conditions (Fig. 2a), so increasing fitness in extreme conditions is reflected in increasing *α* values and decreasing *β* values. So, we model changes in the fundamental niche as the amount of change in *α* and *β*, which is *γn*^*′*^ and −*γn*^*′*^on a logarithmic scale, where *γ* is adaptation rate. This uses the Breeder’s equation. A species with higher adaptation rate evolves its fundamental niche faster in response to environmental deterioration, by evolving higher *α* and lower *β*. We only model speciation events that lead to niche evolution, because speciation events that do not lead to niche evolution, such as allopatric speciation caused by imposition of a physical barrier to interbreeding, do not affect tip niches and so do not influence the posterior probability. As a result, speciation is modelled as a process where an ancestral species’ realized niche splits at a point along an environmental axis and the two descendant species inherit each side of the realized niche. In Figure 2b, the splitting point is determined by parameter *p*, the first *p*-quantiles of the realized niche of the ancestral species. The two descendent species are randomly assigned oppositely signed *p*. The species with positive *p* inherits the right side of the realized niche, so it has higher fitness under the extreme end of the conditions than the ancestral species. The species with negative *p* inherits the left side of the realized niche, so it has lower fitness under these extreme conditions. For both species, the amount of change in *α* and *β* during speciation is *γn*^*′*^ and −*γn*^*′*^ on the logarithmic scale, where *n*^*′*^ is difference in the mean of the realized niche between the new species and the ancestral species. A speciation event can occur at any node in the phylogeny, and occur along any branch when extinction rate is non-zero. When a speciation event occurs along a branch, the descendant species on the left side of the realized niche goes extinct if *p* is positive and the descendant species on the right side goes extinct if *p* is negative. As a result, the speciation event on a branch effectively leads to a punctuated change in the realized niche and the fundamental niche of the ancestral species.

After an adaptation event or a speciation event occurs, there must be a lag between the species’ available niche (the areas it could now occupy given the changed conditions) and the realized niche (the occupied area after selection but before dispersal). Dispersal happens continuously along a section of a branch whose start and end times are defined by a node, an adaptation event, or a speciation event. During dispersal, a species extends its realized niche along the environmental axis towards equilibrium (Fig. 2c) when its realized niche equals the product of its fundamental niche and its available niche. Defining dispersal rate (*d*) as how fast a species reaches the equilibrium, the species realized niche after dispersal is the weighted sum of the realized niche before dispersal and the realized niche at equilibrium, with the latter weighted by the product of dispersal rate and the duration time of the dispersal process. So the faster the dispersal and the longer the duration, the more closely the species realized niche approaches equilibrium.

Defining dispersal in this way, as the ability to occupy areas of favourable conditions, allows us to incorporate dispersal into the model without requiring a spatially explicit model. But we still need to reflect barriers to dispersal in our model, such as emergence of physical barriers to movement, competitive exclusion, or other antagonistic interactions with other species. Since we parameterize the available niche as a continuous distribution, which does not allow any discrete barrier for dispersal, we introduce another parameter *x* to model a barrier by truncating the available niche at *x*. The mechanism for the barrier is not important for our model, so we treat *x* as a nuisance parameter. For example, if during dispersal, a lineage moves to a restricted area, such as an island, then parameter *x* reflects the boundaries on the environmental conditions in that area. For another example, if dispersal follows a speciation event and the location of the barrier (*x*) coincides with the location where a species splits on the environmental axis (*p*) during the speciation event, then parameter *x* reflects competitive exclusion during the speciation event. In this study, we only model dispersal barriers under non-extreme conditions (truncating the side left to *x* in Fig. 2c), because a species distribution under extreme conditions is constrained by the fundamental niche and so little signal for the presence of dispersal barrier under extreme conditions can be detected.

### ENM: inhomogeneous Poisson point process

Above, we described our Niche Evolution Model (NEMo), which models niche evolution along the phylogeny, from the niche at the root to the niches at the tips. NEMo incorporates the change in suitable conditions over time (available niche), the evolution of environmental tolerance (fundamental niche) and the area that the species occupies given dispersal ability and other restrictions such as dispersal barriers (the realised niche). These three aspects of niches at the tips of the phylogeny are then used to calculate *p*(data|niche) using ENM. In principle, any ENM, including process-based ENM, could be used to assess the fit of the inferred niche to the distribution data of a tip species. In this study, we use an inhomogeneous Poisson point process (Renner et al., 2015). The detailed mathematical link between the inhomogeneous Poisson point process and the three aspects of niche is provided in the Supplementary Methods. In some cases, we may have data on the available niche of a tip species, for example, when experts are able to identify areas accessible to a species based on known barriers to dispersal (e.g., Barve et al., 2011). Then *p*(data|niche) has a component for the available niche, which we can calculate as the probability from a Kolmogorov–Smirnov test on the empirical distribution of environmental conditions in the areas known to be accessible to a species, against the available niche of that species as NEMo inferred. In other cases, we may have data on a species fundamental niche, for example, the tolerance range of a species. Then *p*(data|niche) has a component for the fundamental niche, which we can calculate as the probability that the species cannot survive under conditions that are more extreme than the known tolerance limits, given the inferred fundamental niche of the species.

### PCM: reversible-jump model

NEMo includes three biological processes: adaptation, speciation, and dispersal. A phylogenetic comparative method (PCM) is needed to estimate the number and the location of each type of event on the phylogeny. Here we assume adaptation events occur randomly according to a homogeneous Poisson process with rate *r*_*A*_, and speciation events occur randomly according to an inhomogeneous Poisson process with overall rate *r*_*S*_ (Fig. 1). The occurrence of speciation is inhomogeneous because a speciation event may occur at a node or along a branch. Note that we only consider the speciation event that results in niche evolution, which does not necessarily occur at a node. When a speciation event occurs along a branch, it must also be accompanied by an extinction event and the occurrence probability of the extinction event depends on the time when the speciation event occurs. We use reversible jump Markov Chain Monte Carlo (rjMCMC; Green, 1995) to implement the process in a Bayesian framework. Several PCM studies have used rjMCMC to study jumps in evolutionary processes of phenotypic evolution (e.g., Uyeda & Harmon, 2014) and diversification (Rabosky et al., 2014). In our implementation, each jump is an occurrence of a discrete event: an adaptation event or a speciation event. As in these previous studies, our implementation involves several steps (details given in the Supplementary Methods). We implement our method in R (see Data Availability). Given the central role of our niche evolution model in our method, we name the method “NEMo”.

### Simulation study

We performed a simulation study to assess the performance of NEMo given the assumptions that species niches evolve according to our niche evolution model, and processes occur across phylogenies under the Poisson process. In each simulation, we generated species distribution data by 1) simulating phylogenies, 2) randomly placing adaptation and speciation events on the phylogenies, 3) evolving niches along phylogenies to get tip niches, and 4) generating species distribution data from tip niches. Simulation details and choices of parameter values are in Supplementary Methods.

We then use NEMo to reconstruct the history of niche evolution, using only the distribution data generated at the tips and the simulated phylogeny. We tested two aspects of the performance of NEMo: (1) accuracy of estimating the fundamental niche of each tip species, and (2) power to correctly identify the occurrences of adaptation and speciation events on the phylogeny. For the first test, we regressed parameters *α* of each tip species in the simulation against those estimated by NEMo (results for *β* are similar, so we only report *α* here). The slope and *R*^2^ of the regression are reported in Figure 3. We also compared the power of NEMo to a standard ENM (the Poisson point process) to correctly identify tolerant species. For the second test, we calculated power as the proportion of branches with true events in the simulation that are correctly identified by NEMo (see Fig. 3). A branch is identified if it has significantly higher posterior probability of having speciation or adaptation events than expected. We calculated the false positive rate as the proportion of branches with no true events in the simulation that are falsely identified as having events by NEMo (see Fig. 3). Details of these tests are in Supplementary Methods.

**Figure 3.**
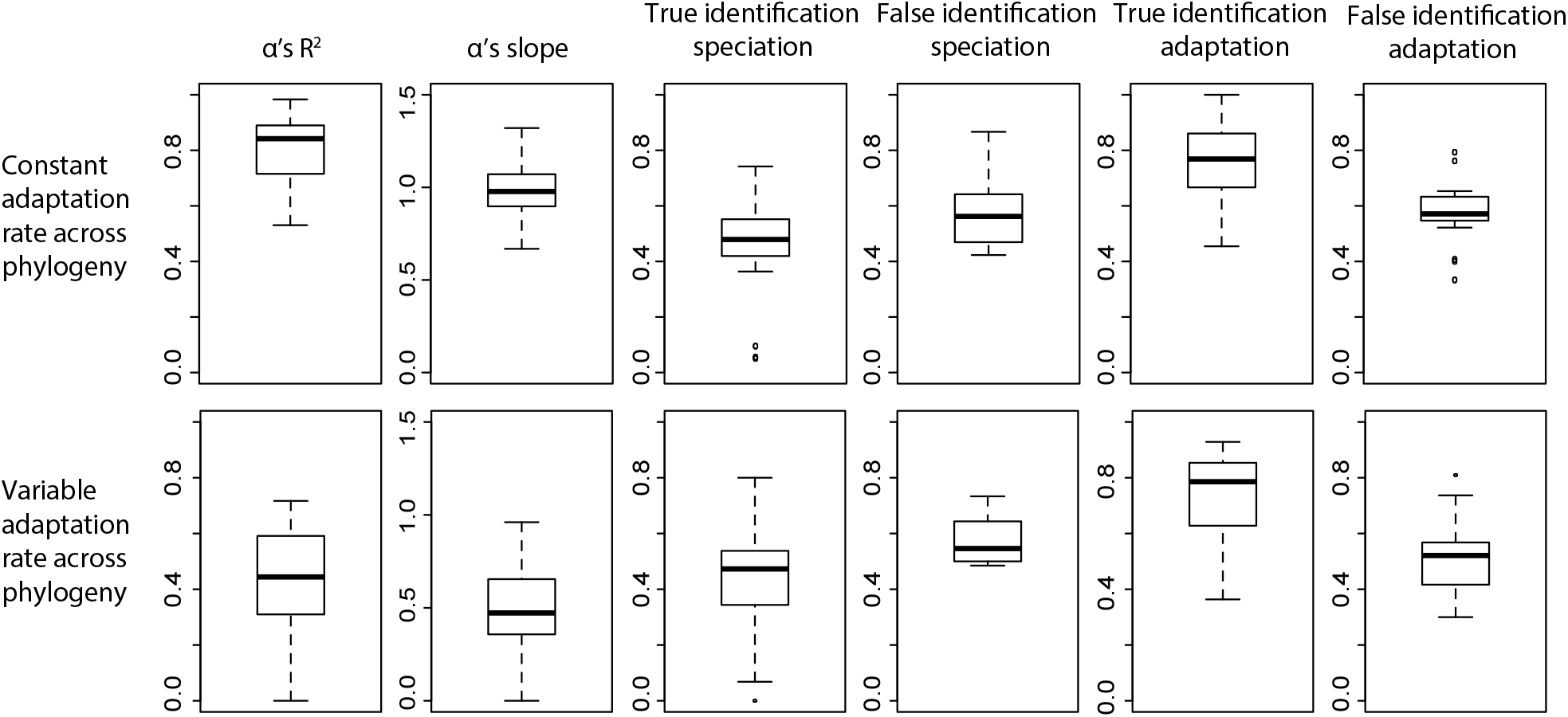
Performance of NEMo on simulated data. Six statistics were used to assess the performance. See text for details on these statistics.

### *Case study: drought- and salt-tolerance in* Acacia

We demonstrated how NEMo can be used to test hypotheses of niche evolution in real datasets, using the example of drought tolerance and salt tolerance in Australian members of the plant genus *Acacia*. Details of the data and the implementation of NEMo in the *Acacia* case study are given in bioRxiv.

We test the utility of NEMo for identifying historical processes of niche evolution, focusing on the evolution of drought tolerance in Australian *Acacia*. This is a useful test case because we can use the climatic history of Australia to generate empirically based *a priori* hypotheses against which to compare our model-based inference from distribution data. The test starts with using NEMo to obtain the posterior samples of the history of niche evolution. From the samples, we plot the histogram of the dates of adaptation events, with the date of each event weighted by the amount of environmental change (parameter *n* in Fig. 1) for that event. Next, we corrected the relative frequency in each bin or date interval of the histogram by the number of branches on the phylogeny during that interval, so that the frequency gives the relative amount of environmental change happened on each branch during the date interval in history, as inferred by NEMo. Last, we compare this distribution of adaptation events to the climatic history of Australia. If NEMo gives reasonable inference on the historical processes of niche evolution, then more environmental change for the adaptation events should be inferred, along the axis of drought tolerance, during periods of aridification than other periods.

We also test the utility of NEMo for inferring the range of conditions under which contemporary species can persist, focusing on salt tolerance in *Acacia*. This is because we can use independently derived information on salt tolerant species to test the adequacy of our inference. We first assemble a list of 12 known salt tolerant species from independent evidence (Table S1). We then use NEMo to obtain the posterior samples of parameter *α* of the fundamental niche of each tip species. From the samples, we calculated the mean of *α* for each tip species. If NEMo gives reasonable estimates for the fundamental niche of each tip species, then the estimated *α* of the 12 known salt-tolerant *Acacia* species should be at the high end of the distribution of values of other *Acacia* species. A Welch’s *t*-test was used to test for the significance in these differences. As a comparison to a standard ENM that does not incorporate niche evolution, we also estimated the ecological niche along the salinity axis of each *Acacia* species from only ENM. Details of the ENM are in Supplementary Methods.

## Results

### Simulation study

Overall, simulations suggested that NEMo performed well under the assumptions of the method, with high accuracy in estimating the fundamental niches of tip species and reasonably high power to identify the occurrence of niche evolution events on the phylogeny. Over all the simulations, the average R squared is above 0.8 and the average slope is around 1, suggesting that the estimates of parameter *α* of the fundamental niches of tip species are highly correlated with their true values in the simulations (Fig. 3). NEMo (power = 0.73±0.19) and the standard ENM (power = 0.73±0.24) have similar power to correctly identify tolerant species. Simulations where the standard ENM outperforms NEMo are when our method has poor MCMC convergence. Simulations where our method outperforms the standard ENM involve many species with narrow realized niches restricted by dispersal barriers.

The power of NEMo to correctly identify branches with adaptation events (true identification of adaptation) is on average 0.8 over simulations, and the power to correctly identify branches with speciation events (true identification of speciation) is on average 0.5 over all simulations (Fig. 3). However, NEMo has a high rate of false identification for both adaptation and speciation events, which is on average 0.6 over all simulations (Fig. 3).

Inferred speciation events in the early diversification history of a species group are more likely to be ‘false’ events (Fig. S1a). Inferred adaptation events with lower amount of environmental change are more likely to be ‘false’ events (Fig. S1b).

An unrealistic assumption of our method is constant adaptation rates across lineages, which is unlikely to be true in the real world, as species’ responses to natural selection depend on many factors. To assess the performance of NEMo when this assumption is violated, we simulated the adaptation rate of each branch on each simulated phylogeny as a random variable. As expected, NEMo has reduced accuracy in estimating the fundamental niche of each tip species under the violation, with the average R squared reduced to 0.5 and the average slope reduced to 0.5 (Fig. 3). However, the violation does not affect the power of our method to identify the occurrence of niche evolution on the phylogeny. The true identification and false identification rates of both adaptation events and speciation events do not differ significantly from those under the simulation scenario of constant adaptation rate (Fig. 3).

### Drought and salt tolerance in Acacia

Here, we report results specific to the predictions we made above to demonstrate the utility of NEMo. But NEMo also infers the process of niche evolution along each branch of the *Acacia* phylogeny, so it reveals new aspects of *Acacia* history. These results and implications are provided in Hua et al (2021 bioRxiv). We used NEMo to reconstruct the history of adaptation to aridity in *Acacia*, using only contemporary observations on living species. The results paint a picture of *Acacia* history that fits with independent historical information from a range of sources. Miller et al. (2013) estimated the age of the common ancestor of extant *Acacia* as 23Ma at the earliest, and since then there have been two key periods of declining temperatures and increasing aridity: a long period from 14-5Ma, followed by a brief warmer and wetter period at the start of the Pliocene (ca. 5Ma), then a second period of aridification from the Late Pliocene (ca. 2.5Ma) and through the Pleistocene (Owen et al., 2017). NEMo identified two peaks in the amount of increasing aridity during *in-situ* adaptation events per branch at 13 Ma and near the present (Fig. 4a), corresponding to the two drying periods at 14-5Ma and after 2.5 Ma, as we expected.

**Figure 4.**
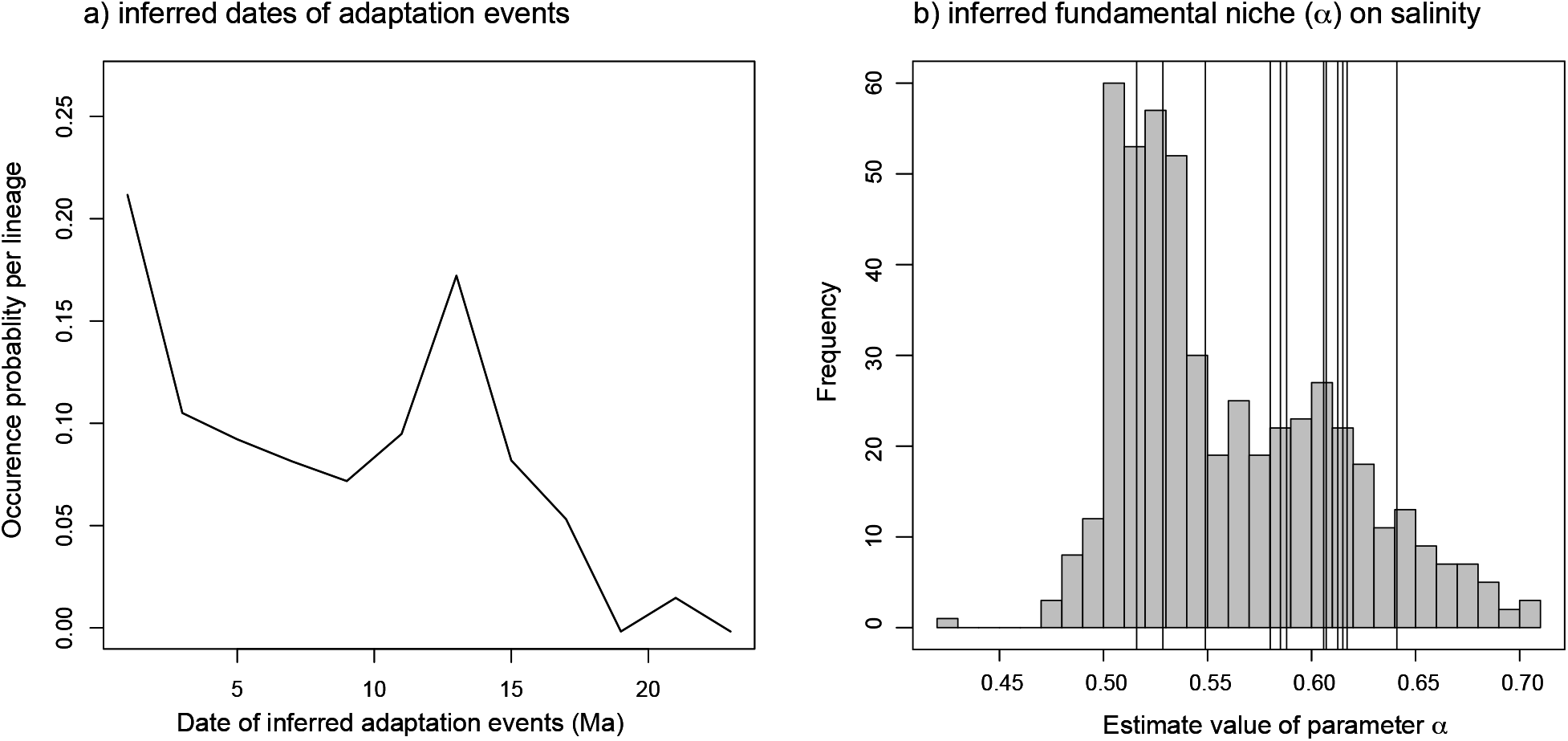
Results for *Acacia* case study. a) Inferred dates of adaptation events in Acacia drought tolerance. The relative frequency of the histogram is weighted by the amount environmental change (*n*) in each adaptation event and is corrected by the number of lineages in the phylogeny during each time interval. b) Inferred fundamental niche with respect to salinity of *Acacia* species. The histogram shows the estimated values of parameter *α* over all the *Acacia* species included in the phylogeny, with larger values indicating greater tolerance to salinity. Each vertical line shows the estimated value of parameter *α* of the 12 known salt-tolerant species.

We also used NEMo to predict which *Acacia* species are highly salt tolerant, based only on their current distribution data. Estimated values of parameter *α* of all the tip species have two modes, with 9 out of the 12 known salt-tolerant species distributed around the mode of higher value (Fig. 4b). The 3 species around the mode of lower value are *A*.*auriculiformis, A*.*sibilans*, and *A*.*harpophylla*. These species belong to clades that are currently distributed in areas with relatively low salinity in Australia, so neither their current distribution nor their relatives’ current distribution provide information on their tolerance range. In general, estimated values of parameter *α* of known salt-tolerant species are significantly higher than the estimated values of the other *Acacia* species (Welch’s *t*=2.48, *p*=0.01; Fig. 4b). In comparison, using only ENM without accounting for shared history of niche evolution among species, the known salt-tolerant *Acacia* species do not have higher fitness under high salinity compared to the other *Acacia* species (Welch’s *t*=0.85, *p*=0.21).

## Discussion

We have presented a new method, the Niche Evolution Model (NEMo), that combines ecological niche models (ENM) and reconstruction of niche evolution within a common statistical framework. The approach we present in this paper models the way that species present-day niches are shaped by evolutionary history and allows us to test historical hypotheses of niche evolution. Using simulations, we show that adding historical processes on the realized niche (area occupied), available niche (suitable conditions) and fundamental niche (range of environmental tolerance) to ENMs results in better predictions, compared to standard ENMs, of fundamental niches from contemporary distribution data, particularly for species with distributions restricted by dispersal barriers. We were also able to better identify salt-tolerant *Acacia* species with this method, using only contemporary distribution data, than the standard ENM. Simulations suggest that NEMo has reasonable power to identify key historical processes of niche evolution on the phylogeny. NEMo captures occurrences of important events in the evolution of drought tolerance in *Acacia* species that are predicted from independent information. In addition, NEMo also has high model flexibility in implementation, making it easily applicable to a wide range of specific questions or case studies in macroevolution and macroecology.

### Assumptions and limitations of NEMo

Unlike previous methods that incorporate evolutionary history into niche models (Smith et al., 2018), NEMo does not use phylogenetic distance as a measure of how much information on a species’ ecological niche can be derived from the distribution data of other species, an approach that assumes strong phylogenetic niche conservatism (Münkemüller et al., 2015). Instead, NEMo explicitly infers events of niche evolution on the phylogeny from species distribution data and lets the data tell us the most likely history of niche evolution along each branch of a phylogeny. As a result, the inferred events of niche evolution along the ancestral lineage of any two species describes the relatedness in their niches. But as with all methods, NEMo has its own assumptions.

One major assumption is that the ecological niche of a species has evolved according to our deterministic process-based model. NEMo includes three fundamental processes of niche evolution –adaptation, speciation and dispersal - but the modelling of each process is highly simplified in order to reduce the number of parameters and limit calculation time. For example, we use one parameter (*p*) to model the process of speciation by splitting the realized niche at a particular point, irrespective of the driving cause of the speciation event. Different drivers of speciation may result in different ways of splitting the realized niche. For example, incipient species under divergent selection may actively move apart from each other to occupy their optimal environments rather than simply cut their realized niche at a particular point. However, even with highly simplified processes of niche evolution, NEMo is already more biologically informative than a pure mathematical model, such as the Brownian motion related models used by most previous PCMs (see review by Revell et al., 2008).

Ideally, we want to model the three processes as joint stochastic processes of both niche evolution and spatial population dynamics, as species niches essentially evolve through changes in the relative abundance of individuals with different tolerance abilities to environmental conditions. In theory, we can model this as a Markovian renewal process (Pyke 1961), which models both the time interval and the outcome of events that change the relative abundance of individuals with different tolerance abilities at each spatial location, including mutation, selection, and dispersal events. Such a model would deal with a large number of states, with each state recording individuals with which tolerance ability in which location has offspring or not. Although we can analytically approximate the long-time behaviour of the model (e.g., Grebenkove and Tupikina, 2018), such a model still requires prior information on how environmental conditions at each spatial location change over time. It is therefore not computationally cost-effective and tractable when we don’t have detailed fossil data to inform past species distributions, or detailed paleoclimate or palynological data to inform spatial variation in past climate and habitats.

Since we often don’t have detailed data on paleo-environments, we need to make another assumption on the occurrence of events of niche evolution. In NEMo, we assume these events occur as a compound Poisson process. Like other applications of the compound Poisson process to molecular evolution and trait evolution, this assumption makes NEMo weakly identifiable (Moore et al., 2016); that is, the model can explain variation in the data equally well either by including lots of events each with a small influence on niche evolution, or by inferring few events with a large influence on niche evolution. This caused the large number of falsely inferred events from the simulation data using NEMo, which occurred in cases where there were speciation events deep in the phylogeny, and adaptation events that involved only small amounts of environmental change were inferred on branches along which no such events occurred. This is not surprising, given that these events have a relatively minor influence on the available niche, so their influence on current species distribution may be overwritten by dispersal, making the inference of these events highly sensitive to the occurrence rate of the compound Poisson process. Similarly, simulations also suggested that inference of true events with a large influence on niche evolution were less sensitive to the occurrence rate of the compound Poisson process, and so our method still gives accurate estimates of the ecological niche of tip species and reliable inferences of the occurrence of major events of niche evolution. As a result, we suggest using narrow priors on the event occurrence rates (*r*_*A*_ and *r*_*S*_) in any empirical application of NEMo, in order to reduce the number false events.

Another challenge associated with low model identifiability is low convergence rate, as MCMC chains can spend long time in local optima (Fig. S3 and S4). Poor convergence can result in poorer performance of NEMo than the standard ENM. Developing more efficient algorithms would help address this problem. For example, we can apply standard ENM to infer tip species’ realized niche first, then treat the mode of the inferred realized niche as a trait and apply standard PCM, such as Lévy process (Landis et al. 2013), to find jumps on the phylogeny that are likely to result in the trait value at the tips. The locations of these jumps give us a good starting distribution of events of niche evolution on the phylogeny, which can then be updated using NEMo.

Two additional assumptions of NEMo are a constant dispersal rate and a constant adaptation rate across the phylogeny. Simulations showed that variation in adaptation rate does not influence the power to infer the history of niche evolution, but it reduces the accuracy to estimate the fundamental niche of tip species. These assumptions can be relaxed, for example, by sampling these rates as random variables in MCMCs. However, whether treating adaptation rate as a random variable can improve the performance of our method requires further investigation. In particular, when both adaptation rate and the number of niche evolution events are free to change on each branch, the model becomes unidentified because the amount of niche evolution depends on the product of adaptation rate and the number of events.

### Extending NEMo and incorporating additional data

The problem of unidentified models also occurs in the area of molecular dating, especially when allowing the molecular evolution rate to change freely in molecular dating (Rannala, 2002). In molecular dating, the solution is to add calibrations on divergence dates (Ronquist et al., 2012). Similarly, we can use paleoclimatic data and fossil distribution data to ‘calibrate’ our combined analysis of niche modelling and niche evolution. Information on paleoclimate is useful for adjusting priors for dates of adaptation events. For example, we know there were three major drying periods in Australia during the Cenozoic – in this paper, we used this as an independent source of historical data against which to test the inferences from NEMo. However, for studies aiming to make inferences about the way species adapt over time, this kind of *a priori* information on changing climate could be used to inform the analysis, for example by increasing the prior probability of the occurrence of adaptation events during the evolution of drought tolerance in these periods.

If we have fossil distribution data, we can extract the paleoclimatic conditions at fossil occurrence locations to inform the realized niche of a higher taxon in the past, such as a genus, as it is often hard to identify fossils to species level and place fossils to a specific branch on the phylogeny. Since NEMo infers ancestral niches at any time along each branch on the phylogeny, we can calculate the realized niche of the higher taxon at a time in the past as the sum of the inferred realized niches of all branches that belong to the taxon at that time. Then, it is straightforward to apply ENM to calculate the likelihood of fossil occurrences given the inferred realized niche of the taxon, which is similar to applying ENM to tip species. ENM has been used in several studies to infer ecological niche of fossilized lineages from fossil occurrences and paleoclimatic modelling (e.g., Meseguer et al., 2015; Chiarenza et al., 2019).

## Conclusion

We present a new method (NEMo) that combines the strengths of environmental niche modelling and a phylogenetic approach to modelling niche evolution over time in order to improve inference of both historical patterns of niche evolution and prediction of current species environmental tolerance limits. Our method makes it possible to combine a range of types of data – including distribution data, phylogeny, fossil dates and locations, physiological information on environmental tolerance, and paleoclimatic modelling – to study niche evolution and estimate species niche. Speciation, adaptation, and dispersal are the three fundamental processes underlying both niche evolution and geographic range evolution. By explicitly modelling the three processes, our method paves a promising path to a holistic understanding of species distribution. The next step is to combine NEMo with methods in historical biogeography to move towards an integrated approach to understanding species’ geographic distribution through the lens of both niche evolution and geographic range evolution.

## Data availability

Source code for the method and for the simulation is available at https://github.com/huaxia1985/NEMo

## Acknowledgements

This paper is dedicated to the memory of John La Salle: we are grateful for his support and encouragement in this and other projects, and we will miss his infectious enthusiasm for biodiversity data and research. We thank Joe Miller, Elisabeth Bui, John La Salle, Rebecca Pirzl, and Lee Belbin for useful comments and discussions. We thank Joe Miller for providing the *Acacia* distribution data. We thank Caroline Chong for her contribution to earlier versions of this project.

## Authors’ Contributions

XH, LB, MC conceived the project and wrote the manuscript. XH developed the models, designed methodology, collected the data, and performed analyses.

## References

Barve, N., Barve, V., Jiménez-Valverde, A., Lira-Noriega, A., Maher, S.P., Peterson, A.T., … Villalobos, F. (2011). The crucial role of the accessible area in ecological niche modelling and species distribution modelling. Ecological Modelling, 222, 1810–1819.

Cang, F.A., Wilson, A.A., & Wiens, J.J. (2016). Climate change is projected to outpace rates of niche change in grasses. Biology Letters, 12, 20160368.

Castro-Insua, A., Gómez-Rodríguez, C., Wiens, J.J., & Baselga, A. (2018). Climatic niche divergence drives patterns of diversification and richness among mammal families. Scientific Reports, 8, 8781.

Chiarenza, A.A., Mannion, P.D., Lunt, D.J., Farnsworth, A., Jones, L.A., Kelland, S.-J., & Allison, P.A. (2019). Ecological niche modelling does not support climatically-driven dinosaur diversity decline before the Cretaceous/Paleogene mass extinction. Nature Communications, 10, 1091.

Dormann, C.F., Schymanski, S.J., Cabral, J., Chuine, I., Graham, C., Hartig, F., … Singer, A. (2012). Correlation and process in species distribution models: bridging a dichotomy. Journal of Biogeography, 39, 2119–2131.

Felsenstein, J. (1985). Phylogenies and the comparative method. The American Naturalist, 12, 1.

González-Orozco, C.E., Pollock, L.J., Thornhill, A.H., Mishler, B.D., Knerr, N., Laffan, S.W., … Bruber, B. (2016). Phylogenetic approaches reveal biodiversity threats under climate change. Nature Climate Change, 6, 1110–1115.

Grebenkov, D.S., & Tupikina, L. (2018). Heterogeneous continuous-time random walks. Physical Review E, 97, 012148.

Green, P.J. (1995). Reversible jump Markov Chain Monte Carlo computation and Bayesian model determination. Biometrika, 82, 711–732.

Hardy, C.R. (2006). Reconstructing ancestral ecologies: challenges and possible solutions. Diversity and Distributions, 12, 7–19.

Hutchinson, G.E. (1957). Concluding remarks. Cold Spring Harbor Symposia, 22, 415–427.

Holt, R.D. (2009). Bringing the Hutchinsonian niche into the 21st century: ecological and evolutionary perspectives. Proceedings of the National Academy of Sciences of the United States of America, 106, 19659–19665.

Kearney, M., & Porter, W. (2009). Mechanistic niche modelling: combining physiological and spatial data to predict species’ ranges. Ecology Letters, 12, 334–350.

Kozak, K., & Wiens, J. (2010). Accelerated rates of climatic-niche evolution underlie rapid species diversification. Ecology Letters, 13, 1378–1389.

Landis, M.J., Schraiber, J.G., & Liang, M. (2013). Phylogenetic analysis using Lévy processes: finding jumps in the evolution of continuous traits. Systematic Biology, 62, 193–204.

Maguire, K.C., Shinneman, D.J., Potter, K.M., & Hipkins, V.D. (2018). Intraspecific niche models for ponderosa pine (Pinus ponderosa) suggest potential variability in population-level response to climate change. Systematic Biology, 67, 965–978.

Meseguer, A.S., Lobo, J.M., Ree, R., Beerling, D.J., & Sanmartin, I. (2015). Integrating fossils, phylogenies, and niche models into biogeography to reveal ancient evolutionary history: the case of Hypericum (Hypericaceae). Systematic Biology, 64, 215–232.

Miller, J.T., Murphy, D.J., Ho, S.Y.W., Cantrill, D.J., & Seigler, D. (2013). Comparative dating of Acacia: combining fossils and multiple phylogenies to infer ages of clades with poor fossil records. Australian Journal of Botany, 61, 436–445.

Mishler, B.D., Knerr, N., González-Orozco, C.E., Thornhill, A.H., Laffan, S.W., & Miller J.T. (2014). Phylogenetic measures of biodiversity and neo- and paleo-endemism in Australian Acacia. Nature Communications, 5, 4473.

Moore, B.R., Höhna, S., May, M.R., Rannala, B., & Huelsenbeck, J.P. (2016). Critically evaluating the theory and performance of Bayesian analysis of macroevolutionary mixtures. Proceedings of the National Academy of Sciences of the United States of America, 113, 9569–9574.

Morales-Castilla, I., Davies, T.J., Pearse, W.D., & Peres-Neto, P. (2017). Combining phylogeny and co-occurrence to improve single species distribution models. Global Ecology and Biogeography, 26, 740–752.

Münkemüller, T., Boucher, F.C., Thuiller, W., & Lavergne, S. (2015). Phylogenetic niche conservatism – common pitfalls and ways forward. Functional Ecology, 29, 627–639.

Owen, C.L., Marshall, D.C., Hill, K.B.R., & Simon, C. (2017). How the aridification of Australia structured the biogeography and influenced the diversification of a large lineage of Australian cicadas. Systematic Biology, 66, 569–589.

Peterson, A., Soberón, J., Pearson, R., Anderson, R., Martínez-Meyer, E., Nakamura, M., & Araújo, M. (2011). Ecological Niches and Geographic Distributions (MPB-49). Princeton, NJ: Princeton University Press.

Pyke, R. (1961). Markov renewal processes with finitely many states. The Annals of Mathematical Statistics, 32,1243–1259.

Rabosky, D. L. (2014). Automatic detection of key innovations, rate shifts, and density-dependence on phylogenetic trees. PLOS ONE, 9, e89543.

Rannala, B. (2002). Identifiability of parameters in MCMC Bayesian inference of phylogeny. Systematic Biology, 51, 754–760.

Renner, I.W., Elith, J., Baddeley, A., Fithian, W., Hastie, T., Phillips, S.J., … Warton D.I. (2015). Point process models for presence-only analysis. Methods in Ecology and Evolution, 6, 366–379.

Revell, L.J., Harmon, L.J., & Collar, D.C. (2008). Phylogenetic signal, evolutionary process, and rate. Systematic Biology, 57, 591–601.

Ronquist, F., Klopfstein, S., Vilhelmsen, L., Schulmeister, S., Murray, D.L., & Rasnitsyn, A.P. (2012). A total-evidence approach to dating with fossils, applied to the early radiation of the Hymenoptera. Systematic Biology, 61, 973–999.

Saupe, E.E., Barve, N., Owens, H.L., Cooper, J.C., Hosner, P.A., & Peterson, A.T. (2018). Reconstructing ecological niche evolution when niches are incompletely characterized. Systematic Biology, 67, 428–438.

Smith, A.B., Godsoe, W., Rodríguez-Sánchez, F., Wang, H., & Warren, D. (2018). Niche estimation above and below the species level. Trends in Ecology & Evolution, 34, 260–273.

Uyeda, J.C., & Harmon, L.J. (2014). A novel Bayesian method for inferring and interpreting the dynamics of adaptive landscapes from phylogenetic comparative data. Systematic Biology, 63, 902–918.

Weber, M.M., Stevens, R.D., Diniz-Filho, J.A., & Grelle, C.E. (2017). Is there a correlation between abundance and environmental suitability derived from ecological niche modelling? A meta-analysis. Ecography, 40, 817–828.

