## Supplementary Methods for "Tracking niche change through time: simultaneous inference of ecological niche evolution and estimation of contemporary niches"

1. Inhomogeneous Poisson point process and the three aspects of niche
2. Reversible-jump model and prior for local parameters
3. Simulation details
4. Case study details

**Figure S1.** The confidence that a branch is correctly identified to contain true speciation events and true adaptation events.

**Figure S2.** MCMC convergence of analyses on simulated data

**Figure S3.** MCMC convergence of analyses on *Acacia* case study

**Supplementary methods:**

*1. Inhomogeneous Poisson point process and the three aspects of niche:*

The Poisson point process models the number of recorded presences of a species per unit area as a Poisson random variable (Renner et al. 2015). The inferred rate of recording a presence of the species in the area depends on the abundance of the species in that area and the sampling intensity in that area (see details in Fithian et al. 2015). Below, we describe how the three aspects of niches are linked to parameters in the ENM, assuming equal sampling intensity across the area (unequal sampling intensity is accounted for in the *Acacia* Case study, with details in bioRxiv).

We can start by considering a dataset where we have presence-only data on distribution: that is, where our knowledge of species niches is informed by records of that species at particular sampling locations for which we can derive relevant environmental data. In brief, the log-likelihood of the presence-only data of each tip species is

$$\ell_P = \sum_{i \in I_{PO}} \log(\lambda_{r_i}) - \mathcal{D} \int \lambda_f \lambda_a ds$$

, where $I_{PO}$ is the set of all the species presence-only locations. In the first term, $\lambda_{r_i}$ is the realized niche, or the rate of recording a presence of the species under the environmental condition at the $i^{\text{th}}$ presence location. The second term is the integral of the product between the fundamental niche ($\lambda_f$) and the available niche ($\lambda_a$), and $\mathcal{D}$ is the geographic area of the available niche. This term describes the overall probability of recording a presence of the species in the species available niche. Often, $\mathcal{D}$ cannot be measured accurately without knowing the dispersal ability of a species. However, it can be used to adjust how informative presence-only locations are for inferring the species fundamental niche. For example, a species with widely scattered presence records has a higher $\mathcal{D}$ value than a species with more spatially concentrated records, so in the species fundamental niche, the same amount of decrease in fitness in conditions not in the presence locations will increase the model likelihood more for the species with higher $\mathcal{D}$ value.

Now we can consider the case where in addition to presence data (indicating points where the species is known to occur), we also have absence data (locations where the species was actively sought but not found). Absence of a species is often inferred from surveys within a local area, such as quadrat sampling. The log-likelihood of the presence-absence data of each tip species is

$$\ell_A = \sum_{i \in I_{PA}} \left[ -\mathrm{y}_i \log\left(1 - e^{-\lambda_{ri}}\right) + (1 - y_i)\lambda_{r_i} \right]$$

, where $I_{PA}$ is the set of all the presence and absence locations of the species, with $\mathrm{y}_i = 1$ if the species is present at the $i^{\text{th}}$ location and $\mathrm{y}_i = 0$ if the species is absent at the $i^{\text{th}}$ location. We don't need to account for the available niche here, because the survey is complete and so wherever the species does not occur, it is assumed to be absent.

To make our model analytically tractable, we converted the continuous environmental axis into bins. The minimum and maximum bins are determined by the minimum and maximum values of environmental conditions in the study area. The three aspects of niche were converted into discrete distributions using the probability that values from these niche distributions fall in the range of each of the bins. For the bin containing the minimum or the maximum value, it is the probability that values from these niche distributions are lower or higher than the minimum or the maximum value.

*2. Reversible-jump model and prior for local parameters:*

The implementation involves six steps: 1) adding a new event to the model, 2) removing an existing event from the model, 3) updating the location of an existing event on the phylogeny, 4) updating parameters specific to that event, 5) updating parameters shared among events, 6) updating the occurrence rate of an event.

In steps 1-2, the acceptance probability of the addition or the removal of an event, or of a proposed model that has one more or one fewer events than the current model, is *min*(1, *LikelihoodRatio*× *PriorRatio* ×*ProposalRatio* ×*Jacobian*). *LikelihoodRatio* is the ratio of the likelihood of the proposed model to the current model. *PriorRatio* is the ratio of the probability, *p*(model|phylogeny), of the proposed model to the current model. *ProposalRatio* is the relative probability of adding or deleting a move to the current model and we followed arguments in Rabosky (2014, equation 6-7) to set *ProposalRatio*. *Jacobian* is included to account for parameter transformation.

To calculate *p*(model|phylogeny), we assume adaptation events occur randomly according to a homogeneous Poisson process with rate $r_A$, and speciation events occur randomly according to an inhomogeneous Poisson process with overall rate $r_S$ (Fig. 1). The occurrence of speciation is inhomogeneous because a speciation event occurring along a branch must be accompanied by an extinction event and the occurrence probability of the extinction event depends on the time $(t)$ when the speciation event occurs, which is $p_E(t) = \frac{\mu(1-e^{-(\lambda-\mu)t})}{\lambda-\mu e^{-(\lambda-\mu)t}}$ (Kendall 1948), where $\lambda$ is speciation rate and $\mu$ is extinction rate. Given that a speciation event occurs, the probability of the event occurring at a node relative to the probability of the event occurring along a branch is the number of nodes in the phylogeny divided by the integral of $\lambda p_E(t)$ over all the branch lengths of the phylogeny. When a speciation event occurs at a node, it has equal probability of occurring at any node in the phylogeny. When a speciation event occurs on a branch, the relative probability that the event occurs on a specific branch depends on the integral of $p_E(t)$ from the starting and the ending times of that branch, and the relative probability of the event to occur at time $t$ in that branch is $p_E(t)$.

We sampled values of each parameter for each event from the same prior distribution as all the other events, so there is no parameter transformation in our implementation and so

*Jacobian*=1. For an adaptation event, parameter $n$ (the number of steps of environmental change) has a discrete prior of equal probability for any integer between one and the number of bins on the environmental axis. For a speciation event, parameter $p$ (the location on the environmental axis where species split) has a discrete prior with equal probability for a sequence from -0.9 to 0.9 by 0.2 increment. For the dispersal process following an event or a node, parameter $x$ (the location of dispersal barrier on the environmental axis) has a discrete prior with equal probability for any integer between 0 and the number of bins on the environmental axis.

In steps 3-4, an existing event was randomly chosen to update. The location of the chosen event on the phylogeny was updated by either shifting the location by a small random quantity that was sampled from a uniform distribution or by moving the event up and down one branch. The frequency of the two ways of proposing a new location is arbitrarily set to 10:1. The proposed location is accepted with probability $min(1, LikelihoodRatio \times PriorRatio)$, where *PriorRatio* is the ratio of the prior probability of the proposed location to the current location of the chosen event. Parameter $x$ and $n$ or $p$ for the chosen event is updated with equal probability of moving the current value to its adjacent lower or higher value in its discrete prior. For example, if the current value for $x$ is 1, the proposed value can be either 0 or 2. The proposed value is accepted with probability $min(1, LikelihoodRatio \times ProposalRatio)$, where *ProposalRatio* is included because the proposal probability is not symmetric: the proposed value can only be higher or lower than the current value, if the current value is at the minimum or maximum bound of the discrete prior.

In steps 5-6, each global parameter was updated, including the mean and the variance of the root available niche, parameter $\alpha$ and $\beta$ of the root fundamental niche, dispersal rate ($d$), adaptation rate ($\gamma$), the occurrence rate of adaptation events ($r_A$), and the overall occurrence rate of speciation events ($r_S$). The mean of the root available niche is updated by adding its current value with a uniform random variable between -$\varepsilon$ and $\varepsilon$, where $\varepsilon$ is a tuning parameter, and the accepted probability is *min*(1,*LikelihoodRatio*). All the other parameters are strictly positive, so they are updated by multiplying the current value with $e^{\eta(U-0.5)}$, where *U* is a uniform random variable between 0 and 1, and $\eta$ is a tuning parameter. The proposed value is accepted with probability *min*(1, *LikelihoodRatio*$\times e^{\eta(U-0.5)}$)).

*3. Simulation details:*

Phylogenies were simulated under a birth-death process using the R package "diversitree" (FitzJohn 2012) with 100 tips, a birth rate of 0.1, and a death rate of 0.03. On each simulated phylogeny, we randomly placed 20 adaptation events with equal probability of placement at any location on the phylogeny, and we placed 20 speciation events with different probabilities of different locations, as described above for *p*(model|phylogeny). In terms of parameter settings, all local parameters were randomly sampled from their corresponding prior distributions. For global parameters, one parameter setting was used for root niches across all simulations, since root niches do not affect how informative species distribution data are to their niches and the history of niche evolution. The root available niche has mean of 1 and standard deviation of 2, and the root fundamental niche has $\alpha$ of 3 and $\beta$ of 2, both along an arbitrary environmental axis that is discretised into 10 bins. Dispersal rate and adaptation rate affect the informativeness of species distribution data. Dispersal rate determines how fast species realized niche reaches the equilibrium set by the fundamental niche and the available niche, so the higher the dispersal rate, the more informative current species distribution is, and so the better the performance of NEMo. To test NEMo under the worst scenario, we set dispersal rate to 1. If our birth rate of 0.1 per million years is reasonable in nature, then this dispersal rate means species realized niche is half way to the equilibrium after 1 million years, which is very low, considering that equilibrium is the commonly accepted assumption in ENMs for non-invasive species (Araújo et al. 2005). We do not have any empirical information on the adaptation rate ($\gamma$) in nature. So we applied two schemes to simulate the best and the worst scenario for the performance of NEMo. The best scenario follows the assumption of a constant adaptation rate across the phylogeny, where $\gamma$ was fixed to 0.03, which is the mean of its prior that was used to ensure a finite likelihood value. The worst scenario highly violates the assumption of a constant adaptation rate across the phylogeny, where $\gamma$ on each branch was randomly drawn from an exponential distribution with mean of 0.03, so many branches having $\gamma$ near zero.

Evolving species niches through the randomly placed events along the branches from the root to each tip according to our process-based model produces the available niche, the realized niche, and the fundamental niche of each tip species. Given these tip niches, we generated three sets of species distribution data from each simulation. One dataset is presence-only data, in which presence-only locations were randomly drawn for each tip species from the species realized niche ($\lambda_r$). The number of presence-only locations is drawn from a Poisson distribution with mean equal to the overall probability of recording a presence of the species in the species available niche. Another dataset is presence and absence data, in which 1000 presence-absence locations were randomly drawn for each tip species with equal probability from each bin of the environmental axis, of which the $i^{\text{th}}$ location has $e^{-\lambda_{ri}}$ probability to be an absence location. The last dataset is background data, in which 1000 locations were randomly drawn for each tip species from the species available niche. The background data is required to perform a standard ENM, so that we can compare the performance of NEMo to the standard ENM. For each simulation, we approximate the posterior distribution of our model by two independent MCMCs for $10^6$ generations, with a thinning interval of 1000 and the first 600,000 generations discarded as burnin. Each chain takes 2.5GHz 1-core processor about three days.

To assess the accuracy of NEMo to estimate the fundamental niche of each tip species, we first estimated, for each simulation, the fundamental niche of each tip species as the average of the values of parameters $\alpha$ and $\beta$ of each tip species in the posterior samples. We then compared these estimated values of parameters $\alpha$ and $\beta$ of each tip species to their corresponding values in the simulation, using linear regression. Last, we summarized the regression results over all the simulations by the slope estimate and the R squared of the regression model. Slope values close to 1 and R squared values close to 1 indicate a high level of accuracy in our estimation of the fundamental niche of tip species.

We also wanted to compare the accuracy in estimating fundamental niche between NEMo and standard ENMs. However, since standard ENMs do not directly estimate fundamental niche but realized niche, we can only compare the power of the two methods to correctly identify tolerant species. For the purposes of this test, we define tolerant species as the top 10 species ranked by their simulated values of parameter $\alpha$ (species with larger $\alpha$ have higher fitness under extreme environmental conditions). Similarly, the top 10 species ranked by the estimated values of parameter $\alpha$ are the tolerant species identified by NEMo. Then, we estimated the realized niche of each tip species in each simulation from only ENM, using the MultispeciesPP R package (Fithian et al. 2015) which implements the same ENM model as used by NEMo. This ENM analysis was applied to the same simulated presence-only and presence-absence data as NEMo and used three customised natural spline bases of the environmental conditions in the simulated background data. Tolerant species identified by the ENM are the top 10 species ranked by the percentage of area under the estimated niche curve that are distributed under extreme environmental conditions. We defined extreme conditions as the condition with environmental value larger than the average over the simulated background data. Last, we compared the percentage of tolerant species identified by the ENM and by our method that are true tolerant species.

To assess the power of NEMo to identify adaptation events, we first calculated for each branch the posterior probability of at least one adaptation event occurring on each branch, by counting the proportion of posterior samples with adaptation events on that branch. We then identified branches that had significantly higher posterior probability than expected if all the branches had equal probability of having adaptation events. Following Shi and Raboksy (2015), a branch had significantly higher posterior probability than expected if its marginal odds ratio was larger than a threshold arbitrarily set to 5. This is essentially hypothesis testing based on Bayes factor for a model with adaptation events on the branch (against a model without adaptation events on the branch), and Bayes factor larger than 5 is generally considered as substantial evidence for the model. Next, we calculated the power of NEMo as the proportion of branches with true adaptation events that passed the test: in other words, how many branches with true adaptation events we have identified. We also calculated the false positive rate of our method as the proportion of branches that passed the test but with no true adaptation events: in other words, how many inferred adaptation events were not true adaptation events.

Last, we wished to find out what kind of branch is more likely to pass the test. We expected that events that occurred more recently, or events with larger influence on niche evolution, are more likely to be correctly identified by our method, so we measured two properties of each branch. For a branch with true adaptation events, one property is the average of the ages of all the adaptation events placed on the branch during simulation. The other property is the sum of the value of parameter $n$ over all the adaptation events placed on the branch during simulation, with a larger value suggesting more niche evolution due to adaptation on that branch. For a branch with no true adaptation events, the same two properties were measured, but they were measured by the events included in the posterior samples instead of the events placed on the branch during simulation. We divided all the branches into groups according to the two properties. For each group, we calculated the ratio of the sum of the posterior probabilities of all the branches with true adaptation events against the sum of the posterior probabilities of all the branches with no true adaptation events. A ratio larger than 1 suggests that, in the group, a branch that passed the test is more likely to contain true adaptation events. We applied the same steps to assess the power of NEMo to identify speciation events, except that the second property was measured by the average of the absolute value of parameter $p$ over all the speciation events placed on the branch during simulation.

*4. Case study details*

For the *Acacia* case study, we used a published dataset that consists of 132,295 presence locations of 508 *Acacia* species (~80% described *Acacia* species in Australia) as well as a published phylogenetic tree that includes the same 508 *Acacia* species (see Mishler et al. 2014). We also assembled species presence-absence survey data from government databases. These presence-absence data were analysed together with the presence locations. We extracted the degree of aridity and salinity at each presence location of each species and each survey site. We also corrected for sampling bias by including two factors that may influence sampling effort at a location: distance to the nearest road and whether it is in a protected area, such as a conservation reserve or national park. More details of the data and the NEMo analysis are available in bioRxiv.

We chose the inhomogeneous Poisson point process (Renner et al. 2015) as the ENM part of NEMo. In order to compare the performance between NEMo and this ENM-only method, we applied both methods to the same *Acacia* species distribution data, the same data on soil electrical conductivity, and the same variables to correct for sampling bias. The ENM-only analysis is conducted using MultispeciesPP R package (Fithian et al. 2015). We used three customised natural spline bases of the soil electrical conductivity to predict a species' ecological niche along salinity axis. We then tested if the 12 known salt-tolerance *Acacia* species (Table S1) have higher predicted fitness, i.e., larger area under their predicted niche curve, under high salinity than the other *Acacia* species, using Welch's *t*-test. We used a different threshold to define high salinity and reported the lowest *p* value of the Welch's *t*-test.

*Reference:*

Araújo, M.B., Pearson, R.G. (2005). Equilibrium of species' distributions with climate. Ecography, 28, 693–695.

Fithian, W., Elith, J., Hastie, T., Keith, D.A. (2015). Bias correction in species distribution models: pooling survey and collection data for multiple species. Methods in Ecology and Evolution, 6, 424–438.

FitzJohn, R.G. 2012. Diversitree: comparative phylogenetic analyses of diversification in R. Methods in Ecology and Evolution, 3, 1084–1092.

Kendall, D.G. (1948). On the generalized ''Birth-and-Death'' process. The Annals of Mathematical Statistics, 19, 1–15.

Mishler, B.D., Knerr, N., González-Orozco, C.E., Thornhill, A.H., Laffan, S.W., Miller, J.T. (2014). Phylogenetic measures of biodiversity and neo- and paleo-endemism in Australian *Acacia*. Nature Communication, 5, 4473.

Rabosky, D. L. (2014). Automatic detection of key innovations, rate shifts, and density-dependence on phylogenetic trees. PLOS ONE, 9, e89543.

Renner, I.W., Elith, J., Baddeley, A., Fithian, W., Hastie, T., Phillips, S.J., Popovic, G., Warton, D.I. (2015). Point process models for presence-only analysis. Methods in Ecology and Evolution, 6, 366–379.

Shi, J.J., Rabosky, D.L. (2015). Speciation dynamics during the global radiation of extant bats. Evolution, 69, 1528–1545.

**Table S1. Known salt-tolerant *Acacia* species.** We identified salt-tolerant *Acacia* species from studies that did experiments on *Acacia* species over a range of salinity conditions. "Salinity" reports the highest salinity that a species was able to tolerate in experimental trials, in deciSiemens per metre (dS/m) or millimoles per litre (mM).

| Species | Salinity | Reference |
|---|---|---|
| *A. ampliceps* | 600mM | 1,7 |
| *A. auriculiformis* | 12-31 dS/m | 2-3 |
| *A. cyclops* | 10-48 dS/m | 4 |
| *A. harpophylla* | 30 dS/m | 5 |
| *A. ligulata* | > 4 dS/m | 6 |
| *A. patagiata* | 10-48 dS/m | 4 |
| *A. redolens* | 10-48 dS/m | 4 |
| *A. salicina* | > 4 dS/m | 6 |
| *A. saligna* | > 4 dS/m | 6 |
| *A. sibilans* | > 4 dS/m | 6 |
| *A. stenophylla* | > 4 dS/m | 6 |
| *A. victoriae* | > 4 dS/m | 6 |

Reference:

1. Theerawitaya, C., Tisarum, R., Samphumphuang, T., Singh, H.P., Cha-Um, S., Kirdmanee, C., Takabe, T. (2015). Physio-biochemical and morphological characters of halophyte legume shrub, *Acacia ampliceps* seedlings in response to salt stress under greenhouse. Frontiers in Plant Science, 6, 630.

2. Miah, M.A. (2013). Salt tolerances of some mainland tree species select as through nursery screening. Pakistan Journal of Biological Sciences, 16, 945−949.

3. Patel, A.D., Jadeja, H., Pandey, A.N. (2010). Effect of salinization of soil on growth, water status and nutrient accumulation in seedlings of *Acacia auriculiformis* (Fabaceae). Journal of Plant Nutrition, 33, 914−932.

4. Craig, G.F., Bell, D.T., Atkins, C.A. (1990). Response to salt and waterlogging stress of ten taxa of Acacia selected from naturally saline areas of Western Australia. Australian Journal of Botany, 38, 619−630.

5. Reichman, S.M., Bellairs, S.M., Mulligan, D.R. (2006). The effects of temperature and salinity on *Acacia harpophylla* (brigalow) (Mimosaceae) germination. Rangeland Journal, 28, 175−178.

6. Aswathappa, N., Marcar, N.E., Thompson, L.A. (1987). Salt tolerance of Australian tropical and subtropical Acacias. In: Turnbull J. (Ed.), Australian Acacias in Developing Countries. ACIAR Proceedings, No. 16, pp. 70−73.

**Figure S1.** The confidence that a branch is correctly identified to contain true speciation events (a) and true adaptation events (b). The confidence is measured by the logarithm of the ratio in the posterior probability of an identified branch with true events to an identified branch with no true events. A positive value (red) suggests that the branch is more likely to be correctly identified, that is the identified branch contains true events. A negative value (blue) suggests that the branch is more likely to be falsely identified, that is the identified branch does not contain true events. The confidence depends on two properties of the branch: how much niche evolution occurred on that branch (x-axis) and how old the events are on the branch (y-axis).

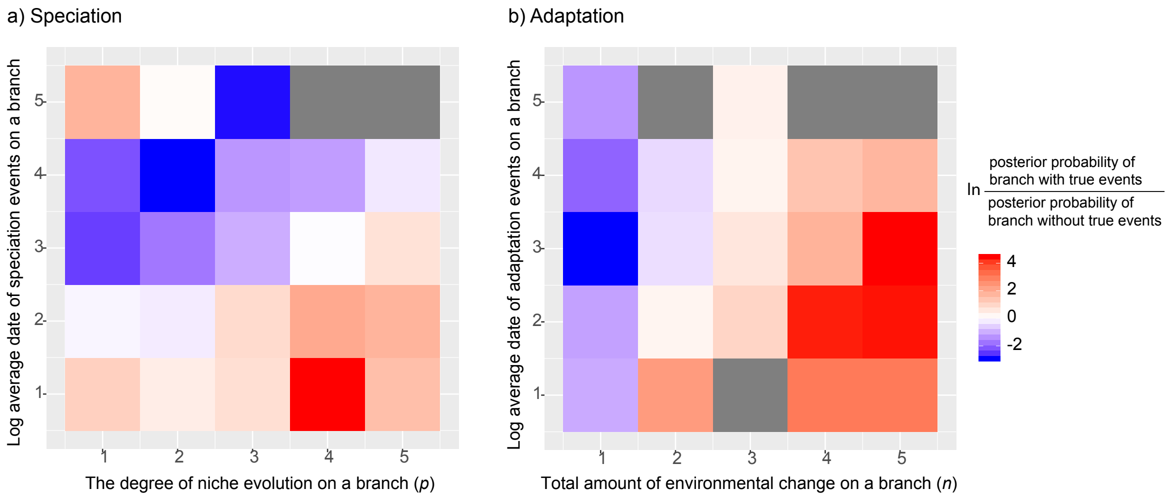

**Figure S2.** MCMC convergence of NEMo on simulated data. a) log posterior probability over all samples including burnin. Different simulations are plotted in different colors. For each simulation, one MCMC chain is in solid line and the other in dashed line. This plot suggests that MCMC chains in most simulations can converge quickly to similar posterior probabilities during the first 100 samples (i.e., $10^5$ generations). b) log posterior probability over samples after burnin. Different simulations are plotted in different colors. Only one MCMC chain is plotted for each simulation to make the plot easier to read. This plot suggests that chains for two simulations do not converge even after $10^6$ generations. These chains probably spent too long at local optima, suggesting that our current rjMCMC algorithm is not efficient at exploring the parameter space, particularly because the acceptance probability of new events of niche evolution depends on the existing events on the phylogeny.

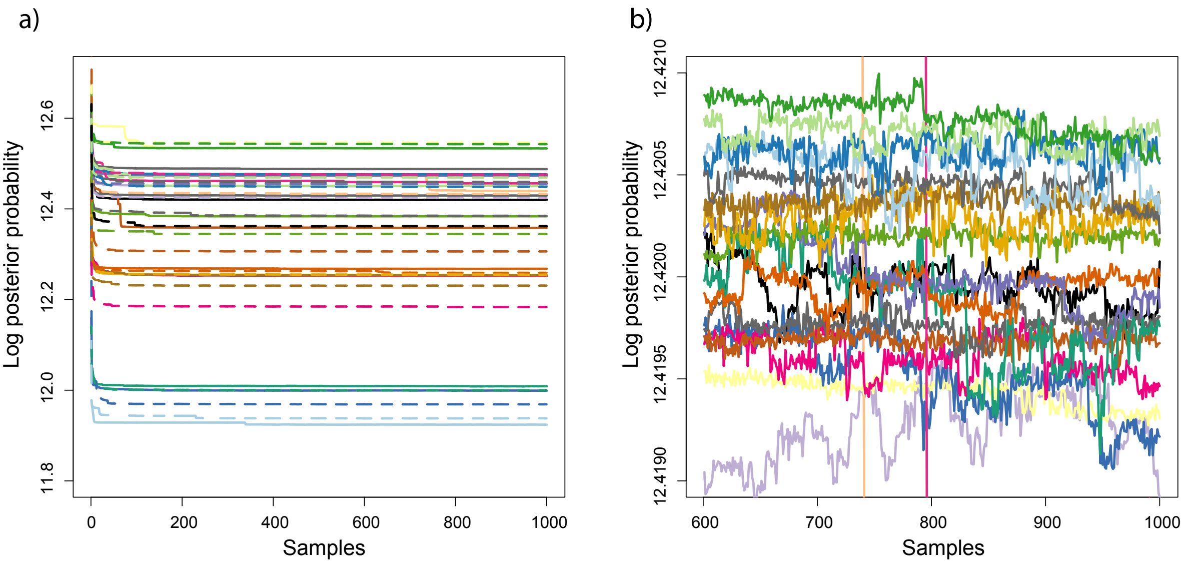

**Figure S3.** MCMC convergence of NEMo on *Acacia* case study. The MCMC chains do not converge to similar posterior probability for the analyses on both drought tolerance (a) and salt tolerance (b) after $10^6$ generations. The low convergence rate is why we used 20 chains, so that we can run multiple chains parallelly to cover as many local optima as allowed by our computing facilities.

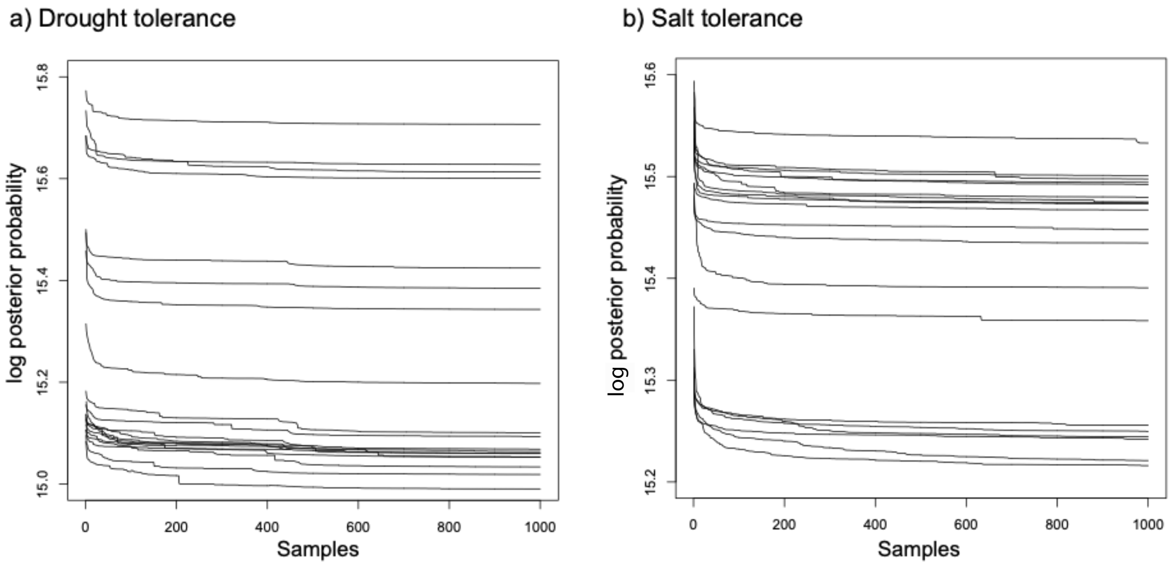
